# Spontaneous and deliberate modes of creativity: Multitask eigen-connectivity analysis captures latent cognitive modes during creative thinking

**DOI:** 10.1101/2020.12.31.425008

**Authors:** Hua Xie, Roger E. Beaty, Sahar Jahanikia, Caleb Geniesse, Neeraj S. Sonalkar, Manish Saggar

## Abstract

Despite substantial progress in the quest of demystifying the brain basis of creativity, several questions remain open. One such issue concerns the relationship between two latent cognitive modes during creative thinking, i.e., deliberate goal-directed cognition and spontaneous thought generation. Although an interplay between deliberate and spontaneous thinking is often indirectly implicated in the creativity literature (e.g., dual-process models), a bottom-up data-driven validation of the cognitive processes associated with creative thinking is still lacking. Here, we attempted to capture the latent modes of creative thinking by utilizing a data-driven approach on a novel continuous multitask paradigm (CMP) that widely sampled a hypothetical two-dimensional cognitive plane of deliberate and spontaneous thinking in a single fMRI session. The CMP consisted of eight task blocks ranging from undirected mind wandering to goal-directed working memory task, while also including two of the widely used creativity tasks, i.e., alternate uses task (AUT) and remote association task (RAT). Using data-driven eigen-connectivity (EC) analysis on the multitask whole-brain functional connectivity (FC) patterns, we embedded the multitask FCs into a low-dimensional latent space. The first two latent components, as revealed by the EC analysis, broadly mapped onto the two cognitive modes of deliberate and spontaneous thinking, respectively. Further, in this low-dimensional space, both creativity tasks were located in the upper right corner of high deliberate and spontaneous thinking (creative cognitive space). Neuroanatomically, the creative cognitive space was represented by not only increased intra-network connectivity within executive control and default mode networks, but also by a higher inter-network coupling between the two. Further, individual differences reflected in the low-dimensional connectivity embeddings were related to differences in deliberate and spontaneous thinking abilities. Altogether, using a continuous multitask paradigm and data-driven approach, we provide direct empirical evidence for the contribution of both deliberate and spontaneous modes of cognition during creative thinking.

## 1. Introduction

It is commonly agreed that creativity refers to the ability to produce work that is both novel and appropriate (Sternberg and Lubart, 1999). As one of the most extraordinary capacities of the human brain, creativity drives the development of our society. From art and design to science and engineering, we often marvel at people’s ingenuity. Given its central role, there has been an ever-growing interest in studying the neural basis of creative cognition. Although initial neuroimaging studies focused on revealing the contribution of individual brain regions to different aspects of creative thinking (Dietrich, 2004; Saggar et al., 2017, 2015), in recent years, this focus has shifted towards examining the interaction between multiple brain regions (as a network) during creative thinking (Beaty et al., 2019, 2017; Maillet et al., 2019; Saggar et al., 2019). However, data-driven evidence is still needed to confirm whether creative thinking depends on a single brain network or an interplay between multiple networks.

As a complex high-level cognitive phenomenon, creativity likely depends on a range of other lower- and higher-order processes, such as perception, working memory, semantic memory, and sustained attention (Dietrich, 2004; Lee and Therriault, 2013; Smeekens and Kane, 2016). Further, an interplay between two latent cognitive modes has been suspected during creative cognition, i.e., modes of spontaneous/implicit thinking and deliberate/explicit thinking. This interplay has been previously referred to as a dual-process model (Barr et al., 2015; Christoff et al., 2016; Dietrich, 2004; Finke, 1996; Sowden et al., 2015). Specifically, previous data suggest that while creative insights are often accompanied by defocused attention through spontaneous thinking (Baird et al., 2012; Eysenck, 1995; Gable et al., 2019; Zabelina et al., 2015), creativity can also stem from methodical problem solving via deliberate thinking (Benedek et al., 2014; Boden, 1998; Frith et al., 2019; Nusbaum and Silvia, 2011).

The interplay between deliberate and spontaneous thinking during creative cognition is hypothesized to correspond to two canonical brain networks: the executive control network (ECN) and the default mode network (DMN), respectively (Beaty et al., 2016, 2015; Ellamil et al., 2012). The ECN is typically elicited by tasks requiring externally driven attention, while the DMN is typically elicited by internally driven cognition. In the context of creativity, the ECN is thought to support goal-directed and strategic cognition required to guide and direct the creative thought process, inhibiting common ideas and strategically searching memory for task-relevant unique solutions (Beaty et al., 2016). The DMN, in contrast, is thought to support the spontaneous generation of candidate ideas from memory and imagination, consistent with its role in episodic/semantic memory retrieval and mental simulation (Buckner et al., 2008). The putative cognitive processes of ECN and DMN broadly map onto dual-process models of creativity that emphasize spontaneous thought and deliberate control (Beaty et al., 2015).

Together, these studies provide insights into the theoretical roles of DMN and ECN in spontaneous and deliberate cognition during creative performance. However, direct data-driven evidence for the involvement of these latent cognitive modes (i.e., deliberate/spontaneous thinking) remains elusive. That is, existing evidence does not unambiguously indicate that ECN and DMN support deliberate and spontaneous cognition during creative performance, while the cognitive roles of ECN and DMN have merely been speculated and inferred based on classic dual-process theories of creativity (e.g., Finke, 1996).

To tackle this issue, here, we adopted a data-driven approach to examine the interplay between spontaneous and deliberate thinking during creative cognition using functional magnetic resonance imaging (fMRI). Specifically, we developed a novel continuous multitask paradigm (CMP) - with seven cognitive task blocks and a resting-state block, in a single fMRI session. Using our CMP, we aimed at sampling a wide range of cognitive processes along a hypothetical two-dimensional plane of deliberate and spontaneous thinking. As shown in **Fig. 1**, we included two established creative tasks (i.e., alternate uses task (AUT) and remote associates task (RAT)), five other non-creative task blocks, and a resting-state block. Based on the theoretical framework by Christoff and colleagues (2016), we hypothesized that the two latent cognitive processes would be differentially recruited by these eight task blocks. For example, tasks such as 2-back working memory that require higher cognitive load would occupy the lower right quadrant, i.e., relying heavily on deliberate thinking while inhibiting spontaneity. Similarly, rest or mind-wandering is likely to recruit spontaneous thinking with minimum deliberate control (top left quadrant). Other non-creative tasks, with a medium level of cognitive load, would reside in the cognitive space between resting-state and working memory. Critically, we hypothesized that creative cognition would require both deliberate and spontaneous thinking and hence occupy the top right quadrant. Leveraging information from a wider variety of cognitive tasks, we aimed to obtain a holistic overview of how creative cognition is related to other cognitive processes. Further, using our CMP we aimed at identifying the latent cognitive axes that may underlie creative cognition. Similar approaches have been recently used to assess the neural correlates of ongoing cognition (Gonzalez-Castillo et al., 2015; Krienen et al., 2014).

**Fig. 1.**
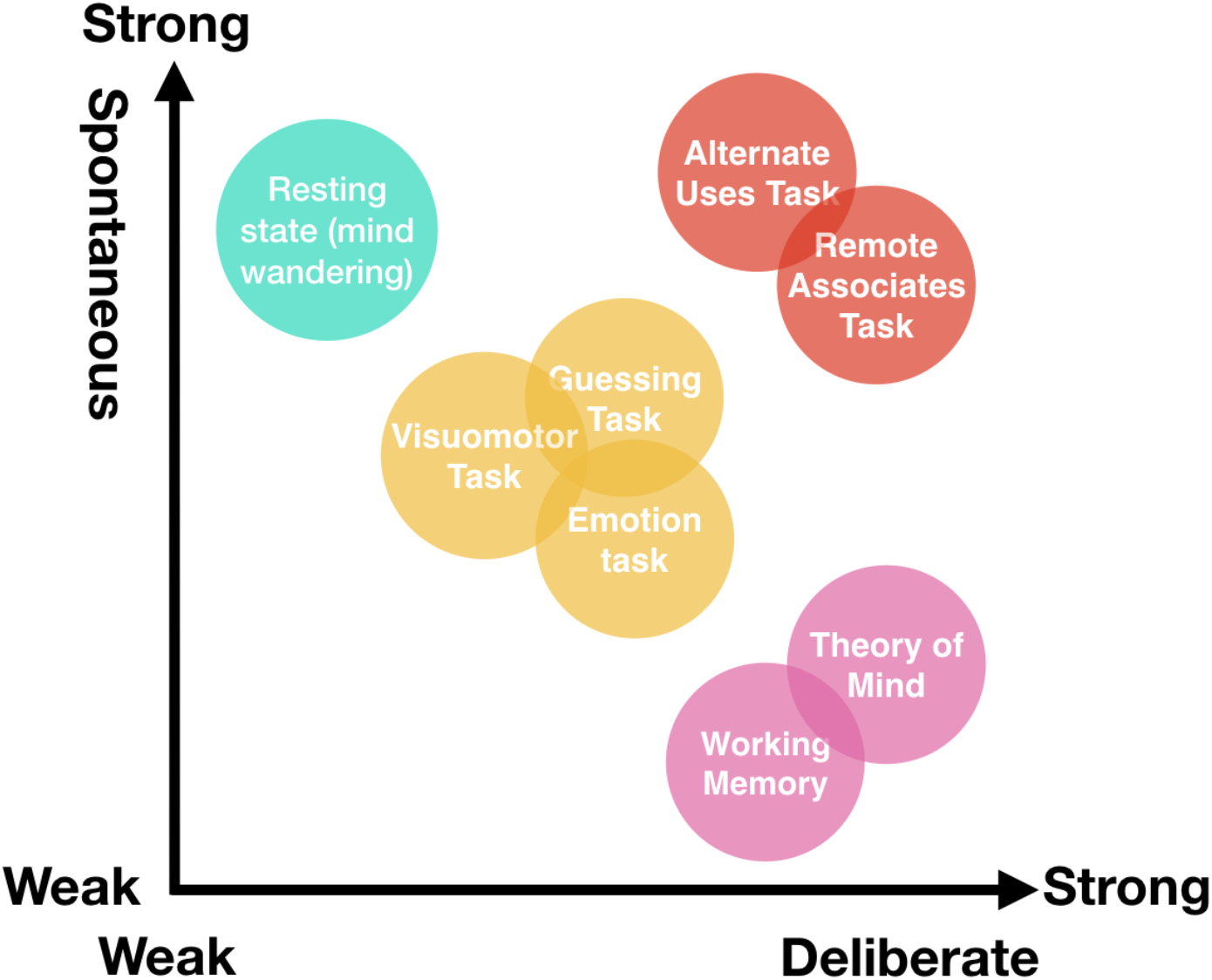
Mapping multitask data to a hypothetical cognitive space with two putatively orthogonal dimensions of deliberate and spontaneous thinking. We hypothesize that tasks such as mind wandering would occupy the upper left quadrant as they are based on spontaneous processing with a minimum amount of deliberate control. In contrast, tasks with high cognitive load (2-back working memory or theory of mind task) would occupy the lower right quadrant as they are based highly on deliberate thinking. Other tasks like emotion classification, guessing, and visuomotor could be mapped in between deliberate and spontaneous thinking. Lastly, we hypothesized that if the creativity tasks (alternate uses and remote association) require both deliberate and spontaneous thinking they should occupy the top right quadrant.

**Fig. 2.**
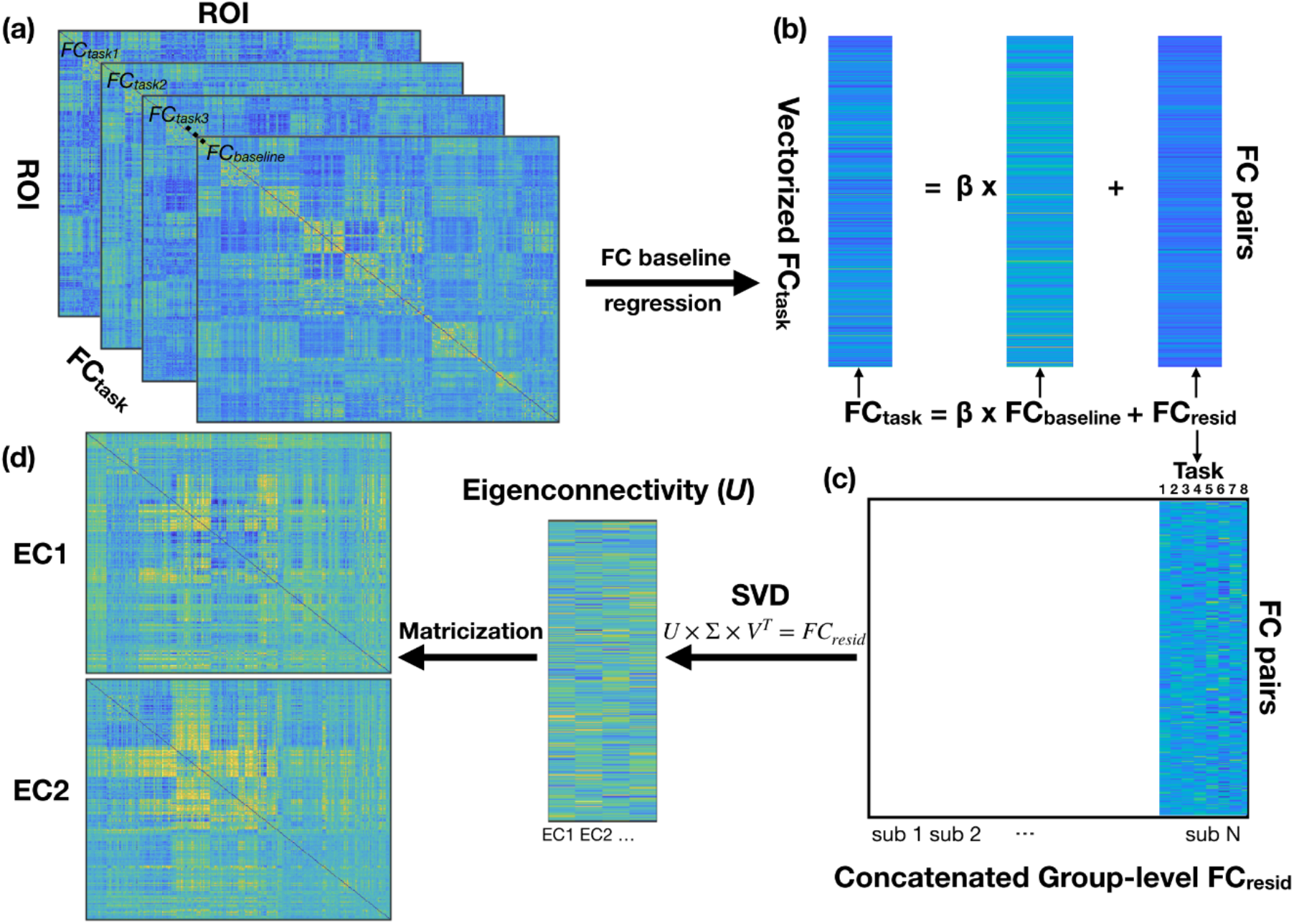
A graphic summary of the EC analysis pipeline. (a) The eight task-specific FC (*FC_task_*) and a baseline-FC (*FC_baseline_*) were computed and then vectorized for each participant. The baseline-FC was computed across the entire scan time. (b) For each participant, the *FC_baseline_* was regressed out from task-specific FCs using linear regression. (c) Baseline-removed FC patterns (*FC_resid_*) were then concatenated across tasks and participants. (d) *FC_resid_* were then submitted to singular value decomposition (SVD). The columns of orthonormal eigenvectors *U* (or equally principal components) were converted to matrix form, termed as eigen-connectivity (EC) patterns.

For a bottom-up data-driven validation of the latent cognitive modes across the eight task blocks, we performed the eigen-connectivity (EC) analysis on the task-related whole-brain functional connectivity (FC) patterns (Leonardi et al., 2013). The EC analysis can reveal the low-dimensional latent embeddings from the multitask FC patterns, i.e., the latent connectivity structures shared across tasks. We aimed to test our hypothetical cognitive space (shown in **Fig. 1**) by examining the low-dimensional embedding revealed by the EC analysis. Further, to estimate the utility of low-dimensional embeddings, we investigated the relationship between individual differences in the latent space embedding and corresponding behavior.

## 2. Methods

### 2.1 Participants

Thirty-two participants (30.4 ± 5.4 years, 13F, 4 left-handed) took part in our study. All participants reported no history of neurological disorder or psychotropic medication, with normal or corrected-to-normal vision. The study was approved by Stanford University’s Institutional Review Board, and all participants gave written consent. Detailed demographic information can be found in Supplemental Table S1.

### 2.2 Neuropsychological assessments

A set of behavioral assessments were conducted outside the MR scanner to measure participants’ creativity and executive function as proxies for spontaneous/deliberate thinking. Below, we briefly introduce these assessments.

#### 2.2.1 Creativity

The Torrance Test of Creative Thinking (TTCT-Figural; Torrance, 1972) is one of the most widely-accepted tests to measure divergent thinking ability in the visual form. This game-like test can engage participants’ spontaneous creativity while being unbiased in terms of race, culture, socio-economic status, gender, and language (Kim, 2006). Participants were given 30 minutes to complete three activities in the TTCT-Figural assessment, i.e., picture construction, picture completion, and repeated figures of lines or circles. The TTCT-Figural assessments were scored by the Scholastics Testing Service, Inc (http://ststesting.com).

#### 2.2.2 Executive function

Participants’ executive function was assessed using the Stroop Color-Word Interference Test (CWIT), a subtest of Delis-Kaplan Executive Function System (D-KEFS; Delis et al., 2001). CWIT consists of four parts: color naming, word reading, inhibition, and inhibition/switching.

### 2.3 Imaging Data

#### 2.3.1 Imaging acquisition

Participants were scanned using a GE 3T Discovery MR750 scanner with a 32-channel Nova Medical head-coil at the Stanford Center for Cognitive and Neurobiological Imaging. Functional scan parameters used are as follows: 1183 volumes, repetition time TR = 0.71 s, echo time TE = 30 ms; flip angle FA = 54°, field of view FOV = 220 × 220 × 144 mm, isotropic voxel size = 2.4 mm, #slices = 60, multiband acceleration factor = 6. High-resolution T1-weighted structural images were also collected with FOV = 190 × 256 × 256 mm, FA = 12°, TE = 2.54 ms, and isotropic voxel size = 0.9 mm.

#### 2.3.2 Continuous multitask paradigm

A novel continuous multitask paradigm (CMP) was conducted over two runs (duration for each run was ~14 min). The CMP included seven cognitive task blocks and a resting state block (Table 1). The cognitive tasks were chosen to sample along the two-dimensional hypothetical plane of deliberate and spontaneous thinking. Each task block lasted 90 s with a 12 s instruction between two task-blocks. A brief summary of the task blocks is provided in Table 1. The CMP was repeated in the second run in a randomized order with different sets of stimuli/questions. Participants were first familiarized with the rules of each task before entering the scanner. For the two creative tasks, alternative uses task (AUT) and remote associates task (RAT), we recorded participants’ answers after the scan, consistent with previous studies (Beaty et al., 2019; Benedek et al., 2019).

**Table 1.**
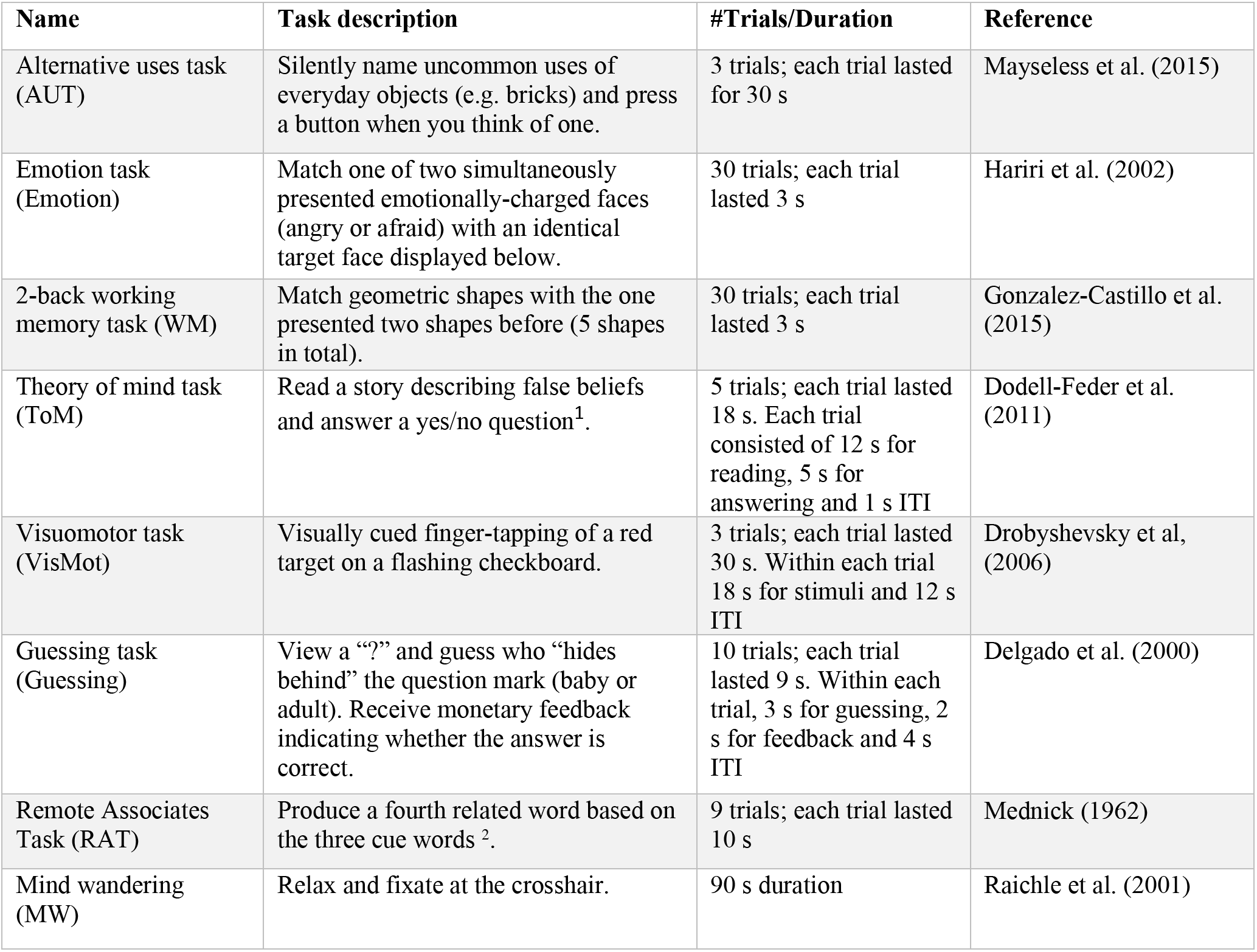
Task batteries included in the continuous multitask paradigm. ITI: inter-trial interval.

#### 2.3.3 Preprocessing

We discarded the first 12 frames of functional data, after which we applied a standardized preprocessing pipeline using fMRIprep (v1.2.1, Esteban et al., 2019). The functional data underwent motion correction, slice timing correction, susceptibility distortion correction, and were normalized to the Montreal Neurological Institute (MNI152) template. Overall, we excluded 7 participants due to technical difficulty (3), poor structural registration (1), excessive motion (1; mean framewise displacement > 0.2mm); and participants’ dropping out or inability to scan (2). All later analysis included 25 participants.

For the remaining 25 participants, we removed nuisance signal by regressing out the physiological noise (white matter and CSF) and motion-related noise using the Volterra expansion of 6 motion parameters and 2 physiological signals (Friston et al., 1996): 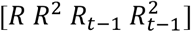. Along with the nuisance signal regression, detrending and temporal filtering between 0.008 and 0.18Hz were also simultaneously performed using AFNI *3dTproject*. Despiking was performed using *3dDespike*, and spatial smoothing was carried out using Gaussian kernel with FWHM = 6 mm. A parcellation with 375 regions of interest (ROIs) were defined based on the parcellation previously used by Shine et al. (2019), which contains 333 cortical parcels from the Gordon atlas (Gordon et al., 2016), 14 subcortical regions from the Harvard–Oxford subcortical atlas (bilateral thalamus, caudate, putamen, ventral striatum, globus pallidus, amygdala, and hippocampus), and 28 cerebellar regions from the SUIT atlas (Diedrichsen et al., 2009) to ensure the whole-brain coverage. After dropping 12 ROIs with fewer than 10 voxels, the time series were extracted from the remaining 363 ROIs by first converting the residual signal to percentage signal change (i.e., voxel intensity was divided by the voxel mean) and then computing the average signal within each ROI. The two functional runs were concatenated, and time points with the framewise displacement greater than 0.5 mm were excluded from further analysis (time points discarded = 1.62%±2.32%).

#### 2.3.4 Estimating regularized functional connectivity (FC)

Sparse graphical models has been increasingly adopted by neuroimaging researchers in recent years (Allen et al., 2014; Rosa et al., 2015; Smith et al., 2011; Xie et al., 2019). Here, we employed graphical LASSO (Friedman et al., 2007) to estimate functional connectivity using the R package ‘glasso’. In short, graphical LASSO encourages a sparse solution of the task-specific precision matrix *Θ* (or inverse covariance matrix) by maximizing the following log-likelihood function *L*_1_

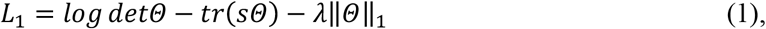

where *det* denotes the matrix determinant; *tr* denotes the matrix trace; s represents the empirical covariance matrix; *λ* is a non-negative regularization parameter provided by users; ||*Θ*||_1_ indicates the L1 penalty on *Θ*.

A zero entry in the precision matrix reflects conditional independence between the signals of two brain regions, after regressing out all other ROI timeseries. A higher *λ* yields a sparser representation at the cost of goodness-of-fit. To achieve a good balance between the sparsity and goodness-of-fit, we tested a range of *λ* (0 - 0.2, step size = 0.02) and found an optimal *λ* (0.06 & 0.08) for each individual that maximizes the following log-likelihood *L*_2_(*λ*)

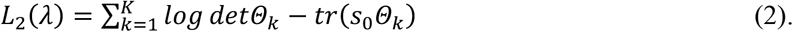

Here, *s*_0_ is the empirical covariance matrix estimated using all the time points, and *K* is the total number of tasks. This objective function was chosen given the expectation that task-specific FCs should be similar across multiple cognitive tasks for a given participant (Finn et al., 2015, also see Supplemental Fig. S1). Upon choosing the optimal regularization parameter, we estimated the regularized covariance matrix and subsequently converted it to regularized whole-brain FC, followed by Fisher-z transformation.

#### 2.3.5 Multitask eigen-connectivity analysis

To delineate the latent cognitive processes sampled by the CMP, we extracted the latent FC structure from the multitask-FC using eigen-connectivity (EC) analysis developed by Leonardi et al. (2013). The EC analysis was originally developed to study time-varying FC dynamics during rest. Briefly, after computing task-specific FC matrices, we first vectorized the upper triangular FC matrices and regressed out the subject-specific baseline-FC to better reveal task-specific FC patterns (Xie et al., 2018a). Here, baseline-FC was characterized as the FC pattern estimated using time points from all eight task blocks for each participant. We then concatenated the residual FC vectors across participants and tasks, resulting in a 65,703 × 200 group-level residual FC matrix (*FC_resid_*) across 8 task blocks and 25 participants. Singular value decomposition (SVD) was applied on the group-level *FC_resid_*.

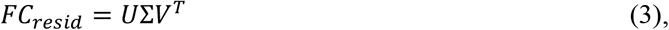

where *U* is a 65,703 × 200 unitary matrix and the columns of *U* are orthonormal eigenvectors; *V* is a 200 × 200 unitary matrix; Σ is a 200 × 200 diagonal matrix of singular values.

The column vectors of *U* were reshaped back into the matrix form (#ROIs × #ROIs). First few columns vectors of *U*, explaining large variance, can be used to define the low-dimensional connectivity-based embedding that is shared across all eight task blocks. The latent embeddings were referred to as EC patterns by Leonardi et al. (2013) when studying dynamic functional connectivity. The EC weights correspond to the projections of these ECs, i.e., columns of *V* multiplied by the singular values of Σ. Given our goal to anchor the cognitive processes into lower dimensions that can be visualized, we focused on the first two ECs that explained the most variance in the group-level *FC_resid_*, as well as the corresponding weights, in order to match the latent cognitive processes of interest.

## 3. Results

### 3.1 Characterizing the latent connectivity dimensions as revealed by EC analysis

We projected vectorized residual task-FCs to a low-dimensional space using the EC analysis. Here, we focused on the first two dimensions/ECs in terms of the variance explained, in accordance with our hypothesis. **Fig. 3a** shows EC patterns for the first two ECs. The strength of intra-network couplings for the first two ECs are shown as line chart in **Fig. 3b**. Each EC pattern can be understood as a latent low-dimensional embedding or spatial mode that captures shared variations across the multitask FCs. On the network level, we observed that the EC1 (shown as upper triangle in **Fig. 3a**) was characterized by a strong intra-network coupling of the ECN (i.e. fronto-parietal network (FPN)) and salience network (SN), which were highest among all networks. On the contrary, the EC2 (shown as lower triangle in **Fig. 3a**) was characterized by highest within network coupling strength of the DMN.

**Fig. 3.**
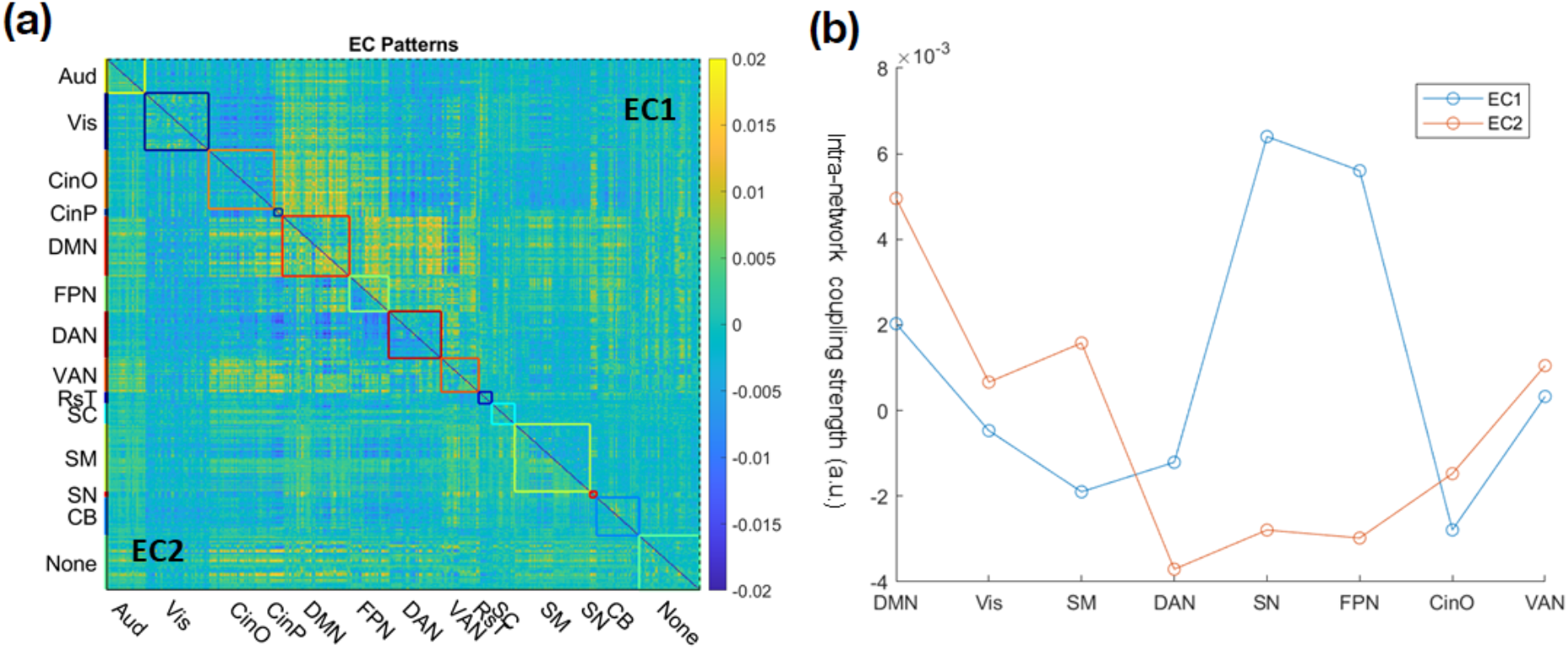
(a) Visualization of the first two eigen-connectivity (EC) patterns with intra-network connectivity highlighted. EC1: the upper-triangle, EC2: lower-triangle. Aud: auditory; Vis: visual; CinO: cingulo-opercular; CinP: cingulo-parietal; DMN: default mode network; FPN: frontal-parietal network; DAN: dorsal attention network; VAN: ventral attention network; RsT: retrosplenial temporal; SM: sensorimotor; SN: salience network; SC: subcortical; CB: cerebellum; None: network not specified. (b) A line plot of the average intra-network coupling strength of major large-scale functional networks.

In sum, the latent space revealed by our multi-task EC analysis suggested two dominant latent dimensions: one dimension for deliberate thinking (characterized by high intra-ECN) and the other for spontaneous thinking (characterized by high intra-DMN).

### 3.2 Embedding tasks into the latent cognitive space

To better examine the relationship between different cognitive tasks with respect to the revealed deliberate and spontaneous EC dimensions, we projected task-FCs through the first two ECs. Noticeably, and as hypothesized, the task-FCs projected into the low-dimensional plane were separable and highly resembled the hypothetical cognitive space (shown as an inset in **Fig. 4a**). Specifically, we observed that the task-FCs associated with two creative tasks (i.e., AUT and RAT) were projected together in the upper right quadrant. The mind wandering (MW) together with the visuomotor (VisMot) task were observed mostly in the upper left quadrant, as both required minimum deliberate control^3^. Working memory (WM) and theory of mind (ToM) tasks were also projected together to the lower right quadrant. These two tasks were arguably among the most cognitively demanding tasks while requiring very limited spontaneous thinking. Further, based on our hypothesis, grouping the tasks into four types: deliberate (WM and ToM), spontaneous (MW), moderate (Emotion, Guessing, and VisMot), and creative (AUT and RAT), revealed that the projection of creativity tasks on EC1 resembles deliberate processing and their projection on EC2 resembles that of spontaneous processing (**Fig. 4b-c**). Altogether, providing data-driven evidence that creative cognition does require an interplay between both deliberate (EC1) and spontaneous thinking (EC2).

**Fig. 4.**
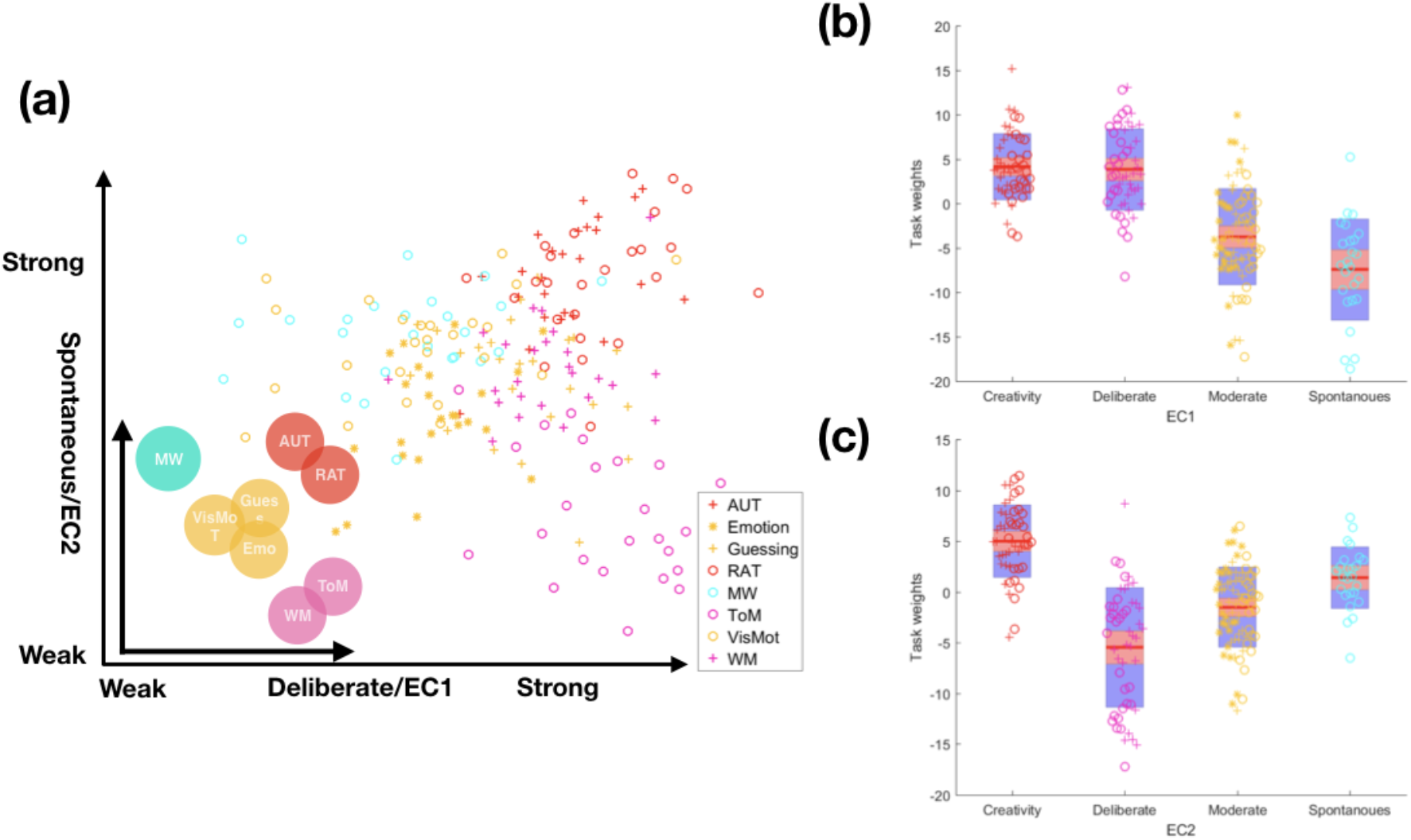
(a) Low-dimensional projection of task-FCs with the first two EC components, color-coded based on task labels. Each symbol represents a projection of a task-FC, for a total of 200 symbols (25 per participant over 8 tasks). Inset: the hypothetical cognitive space spanning across two putative cognitive axes, i.e., deliberate thinking (EC1) and spontaneous thinking (EC2). (b-c) Graphical summary of low-dimensional projections along with the EC1 and EC2, grouped based on the hypothesis. Creativity tasks: AUT and RAT; deliberate tasks: WM and ToM; spontaneous task: MW; tasks requiring moderate level of spontaneous and deliberate thinking: Emotion, Guessing, and VisMot.

Additionally, we showed EC3 to EC10 and the low-dimensional projection using the first 3 ECs in the Supplemental Materials (**Fig. S2**). We also quantitatively evaluated the task separability of weights of all 200 ECs using one-way ANOVA given the task labels. We found ECs beyond the first two can inform us of underlying tasks, where EC weights from one task were significantly different from the rest (FDR-corrected *p* < 0.05, **Fig. S3**).

### 3.3 Revealing the functional architecture of creative cognition

To further understand the functional architecture of the creative cognition space, we examined the aggregated functional connectome across deliberate (EC1) and spontaneous (EC2) latent dimensions. We hypothesized that given the observation that creative tasks were both embedded in the top right corner of the EC latent space, suggesting an interplay between both deliberate (EC1) and spontaneous (EC2) modes, a better understanding of the functional architecture of creative cognitive space can be acquired by examining the aggregated connectivity pattern of EC1 and EC2. The aggregated pattern of EC1 and EC2, through numerical addition, is shown in **Fig. 5a**. Besides the expected enhanced intra-network coupling within default mode and fronto-parietal networks, we also observed high inter-network coupling between default mode and fronto-parietal, cingulo-opercular, and cingulo-parietal networks.

**Fig. 5.**
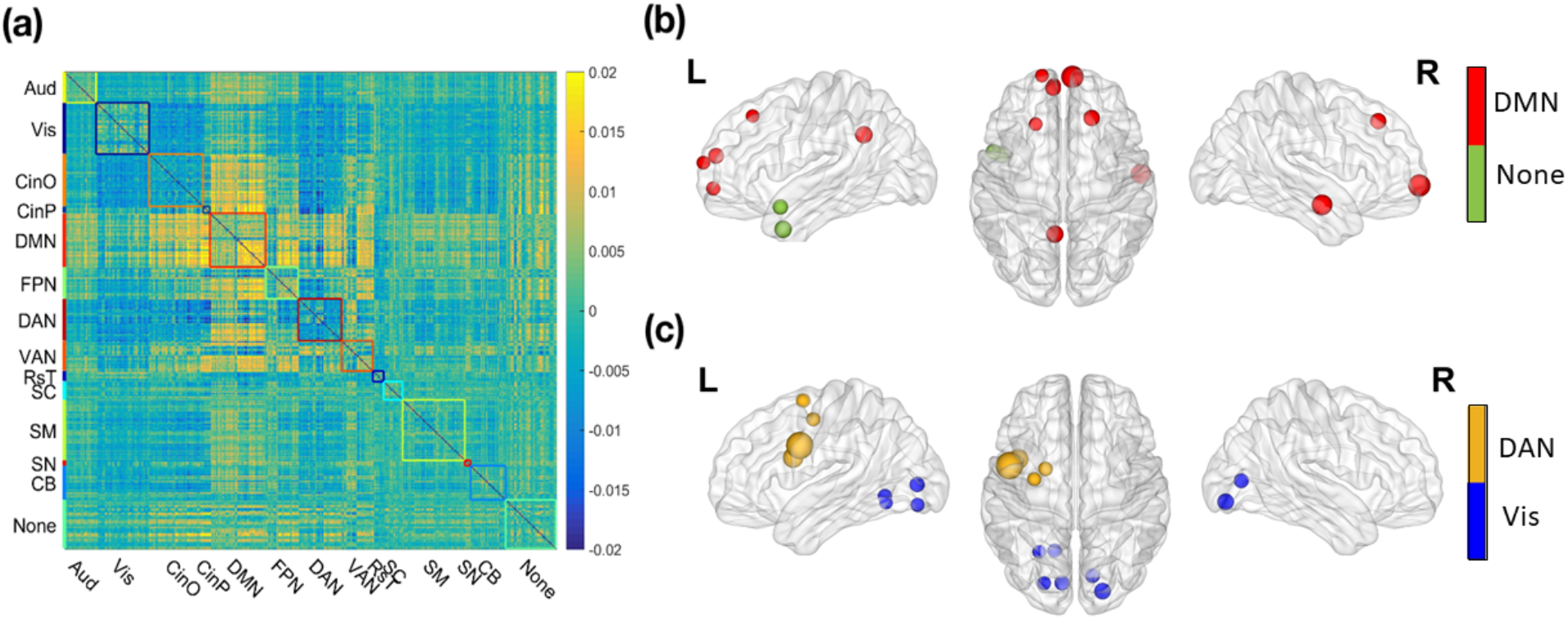
(a) The aggregated EC combining EC1 and EC2. (b) Ten ROIs with the most positive node strength. (c) Ten ROIs with the most negative node strength. The node size is proportional to the node strength, and the network label of each ROI is color-coded.

In an attempt to understand the large-scale network architecture underlying the aggregated EC pattern, we computed the node strength, and for visualization purposes, we showed the ten ROIs with the highest/lowest node strength in **Fig. 5(b)&(c).** In terms of hubs with the positive node strength (i.e., highly coupled regions), the majority was found in the brain regions that form the DMN as well as some in the anterior temporal lobe. The ROI with the highest positive node strength was found to be the right medial prefrontal cortex (mPFC, MNI coordinate: 4.8 65.1 -7.1) of the DMN. On the other hand, hubs with the largest negative node strength (i.e., regions decoupled from other regions) were found in the visual network and the DAN (left lateralized). The ROI with the highest negative node strength was found in the left inferior frontal gyrus (IFG, MNI coordinate: -45.2 2.7 32.4) within the DAN.

### 3.4 Examining whether individual differences in embedding can predict behavior

Individual differences were characterized in terms of EC-based latent-space embedding. We hypothesized that the observed individual differences in the latent-space embedding could be associated with individual differences in behavior. We limited this analysis to the two creativity tasks only. Specifically, the individual differences in the weights of deliberate dimension (EC1) could be related to deliberate thinking ability, while the variability in the weights of spontaneous dimension (EC2) could be related to spontaneous thinking ability. The behavioral correlates of deliberate and spontaneous thinking were computed as follows. We used the behavioral performance on the color-word interference task (CWIT) as a proxy of participants’ deliberate thinking ability. To operationalize behavioral performance of spontaneous thinking, we regressed the CWIT score from the Torrance Test of Creative Thinking task score (TTCT-F). Hence, by removing the variance associated with deliberate thinking from the creativity score, we attempted to use the residuals as a proxy for spontaneous thinking.

After controlling for age, handedness, and gender, we found that the weights of EC1 during the AUT were significantly positively correlated with deliberate thinking score (r = 0.45, *p* = 0.030), and EC2 during the AUT was significantly positively correlated with spontaneous thinking score (r = 0.43, *p* = 0.038), as shown in Fig. 6. We did not observe any significant brain-behavior relationship using EC weights for the second creativity task (RAT).

**Fig. 6.**
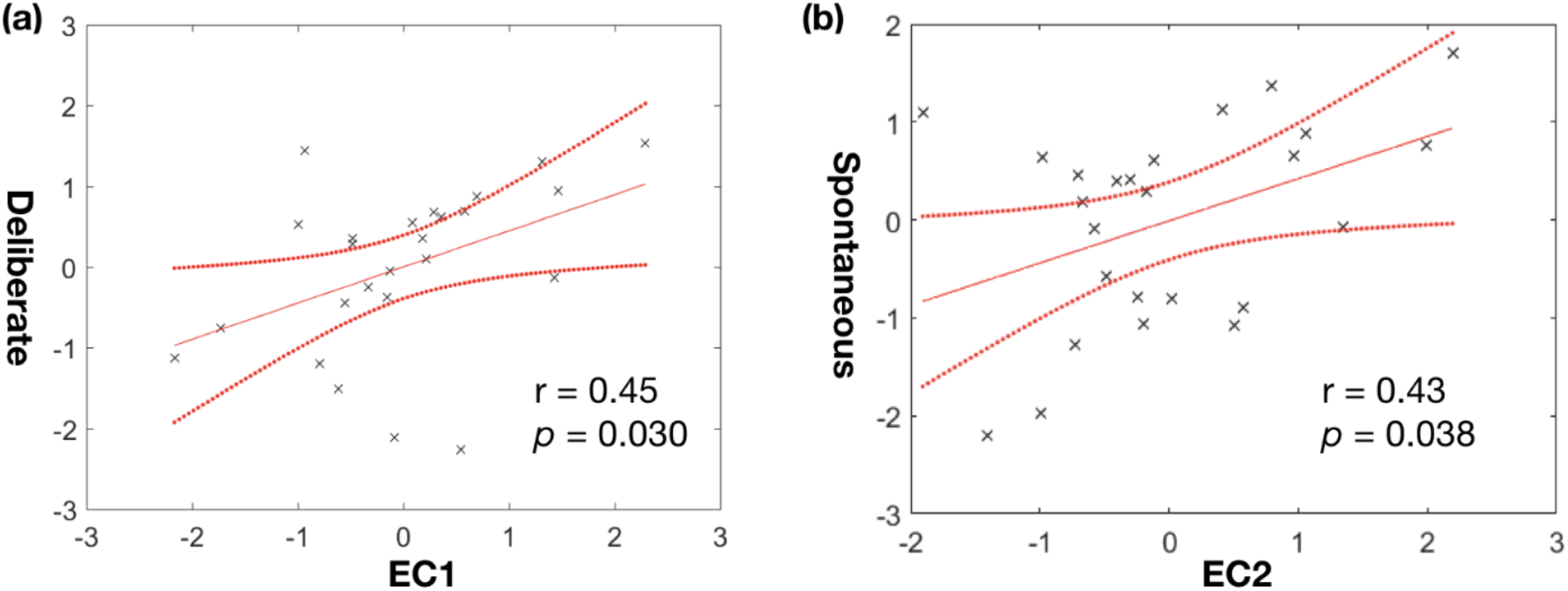
Brain-behavior relationship using EC weights during AUT. The EC weights and behavioral scores are z-scored. (a) The scatterplot of EC1 weights vs deliberate thinking score; (b) EC2 weights vs spontaneous thinking scores. Dotted lines represent 95% confidence intervals.

## 4. Discussion

Human creativity is a vast construct, seemingly intractable to scientific inquiry, partially due to its multifaceted nature (Jung, 2013). It has been long suspected that creative cognition is supported by two latent cognitive modes (i.e., deliberate and spontaneous modes of thinking). However, the neural evidence for the contribution of spontaneous and deliberate cognition in creativity has been indirect and inconsistent (Mok, 2014). Here, to validate the involvement of these latent cognitive modes in creative thinking and to identify their neural substrates, we adopted a data-driven approach and sampled across a wide range of cognitive space using an 8-task continuous multitask paradigm (CMP). We hypothesized that by sampling a wider cognitive space we will better understand how creative cognition is related to other lower- and higher-order cognitive processes.

Since creative cognition does not seem to be confined to any localized brain region (Dietrich and Kanso, 2010), we decided to focus on examining the large-scale network architecture using whole-brain functional connectivity (FC). We first computed the task-FCs and then extracted latent connectivity patterns across all tasks using eigen-connectivity (EC) analysis (Leonardi et al., 2013). The first two latent dimensions were observed to represent the deliberate and spontaneous modes of thinking, respectively. When the task-FCs were embedded into a 2-dimensional latent space of deliberate/spontaneous thinking, we observed creativity tasks to be embedded in the region with both strong deliberate and spontaneous thinking. The embeddings of other tasks also followed as expected. For example, the cognitively demanding tasks such as the theory of mind and n-back working memory appeared to tax deliberate thinking heavily yet requiring little spontaneous thinking. On the contrary, resting state (mind wandering) and visuo-motor task were embedded higher on the spontaneous mode of thinking. Further, the individual differences in EC weights were observed to be related to behavioral differences in the ability of deliberate and spontaneous cognition during a creative task. Altogether, our findings demonstrate the potential of using a data-driven approach to pool information across multiple cognitive processes in order to extract latent cognitive dimensions associated with creative cognition.

Early research on creative cognition focused on isolating specific brain regions associated with creative performance. Although domain-specific assessment of creative cognition proved somewhat successful in teasing out regions specific to each domain, e.g., musical (Limb and Braun, 2008), verbal (Bechtereva et al., 2004), and figural (Ellamil et al., 2012; Saggar et al., 2017, 2015), the domain-generic assessment of creativity revealed a large variance in findings across studies (Boccia et al., 2015). Recently, researchers have shifted gear towards studying the whole-brain functional architecture related to creative cognition. These network-based studies have highlighted a putative role of the default mode network (DMN) and executive control network (ECN) during creative thinking (Beaty et al., 2015; Zhu et al., 2017). In general, while the DMN has been suggested to support spontaneous cognition, such as mind-wandering, introspection, autobiographical memory, and mentalization (Raichle, 2015), the ECN (operationalized as the frontal-parietal network (FPN)), is commonly considered as a key player in deliberate, goal-directed cognition. With regards to creative thinking, the current consensus is that an interplay between deliberate (ECN) and spontaneous thinking (DMN) is required for creative cognition. However, no data-driven validation exists regarding how this interplay facilitates creativity.

To address this issue, here, we used a continuous multi-task fMRI paradigm consisted of a wide range of cognitive tasks including creativity, and explored the latent dimensions using eigen-connectivity analysis. Interestingly, the first two latent dimensions were mapped onto deliberate (FPN-dominated intra-network coupling) and spontaneous (DMN-dominated intra-network coupling) axes. We also examined the extent of inter-network coupling for each latent dimension. For the deliberate axis, i.e., EC1, we observed greater inter-network connectivity between DMN and task-positive networks (including FPN, dorsal attention network (DAN), and cingulo-opercular network (CinO)). Our observation coincided with an earlier finding of increased DMN connectivity with task-promoting regions across six tasks regardless of task-associated activation (Amanda Elton and Wei Gao, 2015). For the spontaneous axis, i.e., EC2, we observed reduced intra-network coupling of the FPN as well as stronger within-network connectivity in the DMN. The weakened within-FPN coupling might allow for flexible reconfiguration during spontaneous thinking, which has been shown to positively correlate with creativity across the visual and verbal domains (Zhu et al., 2017). Moreover, an overall decoupling was observed for DAN, possibly reflecting down-regulated top-down attention modulation (Zabelina and Andrews-Hanna, 2016). Lastly, as creativity required both cognitive modes (EC1 and EC2), we aggregated first two EC patterns and revealed strengthened within-network coupling in the DMN and FPN, as well as an overall increase in inter-network connectivity between the two. Overall, our findings extend network neuroscience research on creative cognition by identifying patterns of intra- and inter-network connectivity associated with latent cognitive modes during creative task performance.

To pinpoint the key regions in the aggregated EC pattern underlying creative cognition, we examined the regions with the highest absolute node strength. The regions with the highest positive functional coupling were found in the DMN, such as mPFC, angular gyrus (AG), and posterior cingulate cortex (PCC), as well as regions in the anterior temporal lobe. The involvement of DMN in creative cognition has been well-documented. For example, higher creativity has been associated with increased FC between the mPFC and the PCC (Takeuchi et al., 2012). A lesion study found that lesions in the mPFC were associated with impaired originality (Shamay-Tsoory et al., 2011). Moreover, using connectome-based predictive modeling (Rosenberg et al., 2015), a recent study found regions in the DMN were among the top contributors to the so-called “high-creative network”, i.e., the network where FC strength positively predicted creativity scores (Beaty et al., 2018). Moreover, the anterior temporal lobe (or temporal pole) has an important role in many cognitive processes, including creative cognition, theory of mind, emotion processing, and semantic processing (Wong and Gallate, 2012). In short, our findings suggested that whole-brain integration of regions in DMN plays a pivotal role in creative cognition.

As for the ROIs with the greatest decrease in the connectivity of the aggregated EC, the majority were found in the DAN and visual network. The decoupling of the visual network is consistent with past work linking deactivation of the visual cortex to the suppression of external stimuli during creative thinking (Benedek et al., 2016; Ritter et al., 2018). On the other hand, as it is well-known that DAN is responsible for external attention (Maillet et al., 2019), decoupling of DAN may also signal loosened top-down attention to external stimuli, potentially allowing for allocating more cognitive resources toward an introspective stream of consciousness (Zabelina and Andrews-Hanna, 2016). Interestingly, we observed left-lateralized decoupling for the DAN. This left-over-right decoupling pattern in DAN mirrors lesion studies linking left hemisphere lesions to increases in creativity (Seeley et al., 2008; Shamay-Tsoory et al., 2011; c.f. Chen et al., 2019). It has been suggested that, under an inhibitory mechanism, the right hemisphere’s predominance in creative cognition may be inhibited by the left hemisphere in typical people, while such inhibition is weakened after damages to the left hemisphere, thus boosting creativity (Huang et al., 2013). In our case, the decoupled left-lateralized DAN (especially L IFG and surrounding ROIs) could be linked to the release of inhibition of the left hemisphere in a similar fashion that facilitates creativity, although the lateralized involvement may depend on the creativity domain (Chen et al., 2019). Moreover, Lotze and colleagues (2014) also noted that a reduced left- and inter-hemispheric connectivity of language areas, namely the left posterior area BA44 (left IFG), may lead to a more spontaneous and less constraining cognition. This is consistent with our observation that the left IFG showed the greatest decoupling in the aggregated EC pattern. Taken together, down-regulation in regions responsible for top–down externally-directed attentional control in the left prefrontal cortex (e.g. left IFG) appears to be a key neural feature for both creative cognition and related spontaneous cognitive processes, such as mind wandering (Christoff et al., 2009; Julia W. Y. Kam et al., 2013).

To sum up, our work sheds new light on the complex interaction between DMN and FPN during creative cognition. Our data-driven approach suggests these typically opposing networks may indeed cooperate during creative cognition as revealed by our EC analysis. Furthermore, decoupling of key regions in DAN and visual networks may also correspond to the shielding of internally directed attention from the external environment during creative thinking (Maillet et al., 2019), further facilitating creative cognition.

### Limitations and future directions

There are some methodological limitations associated with our study. The first issue concerns the relatively small sample size (N = 25), which limited our statistical power in the brain-behavior analysis. Future studies with more participants are needed to further validate our findings, as a recent study suggests that it may require a consortium-level sample size to obtain a reproducible brain-behavior relationship (Marek et al., 2020). Second, previous studies have shown the inter-subject differences in FC patterns are dominated by stable individual differences other than transient cognitive/task modulation (Finn et al., 2015; Gratton et al., 2018; Xie et al., 2018a). We circumvented this issue by removing individual baseline-FCs (i.e., FC fingerprints). However, this is a rather simplified means of removing individual differences by assuming a linear relationship between task-specific and subject-specific FC patterns. Moreover, EC analysis assumes linearity, therefore, it can only capture linear relations among connectivity pairs. Future studies can consider nonlinear decomposition methods such as general principal component analysis (Vidal et al., 2005) and geometry-aware principal component analysis (Harandi et al., 2018). These nonlinear methods could help us more efficiently explore the nonlinear relationships between task-FCs. Third, given our goal of anchoring latent deliberate and spontaneous thinking during creative cognition, we narrowed our focus to the first two EC components. Our choice was partially justified by linking EC weights with behavioral data (Fig. 6), and matching EC patterns with previous neuroimaging findings in creative research. However, there is no doubt that the EC patterns beyond the first two could be meaningful, as many higher-order ECs also explained significantly more variance than those from the surrogate data and provided some task separability (Supplemental Fig. S4). Indeed, some interesting work has been conducted using EC analysis on the multitask data from Human Connectome Project, which looked at higher-order EC components to better identify individuals and tasks (Abbas et al., 2020; Amico and Goñi, 2018). However, as is often the case with any latent factor analysis, increasing the number of latent components/factors comes at the cost of interpretability. We believe that for our specific question, limiting our focus to the first two ECs was a reasonable trade-off. Future work could also look at a different set of tasks with different latent cognitive dimensions. For example, using different forms of creativity (e.g. figural, verbal, and musical) or parametrically modulating the cognitive load (1, 2, 3-back WM), we can delineate the potential confounding factors (e.g., creativity domains and task difficulties). Lastly, the rich spatiotemporal dynamics of the brain remain untapped in this study. Future work can also investigate the time-varying FC during task performance (Gonzalez-Castillo and Bandettini, 2018; Vergara et al., 2019; Xie et al., 2018b) as well as instantaneous activation patterns using Topological Data Analysis (TDA; Geniesse et al., 2019; Saggar et al., 2018). Additionally, our analysis also assumed that the functional parcellation remained unchanged despite the changing cognitive demands, which is subject to future evaluation (Salehi et al., 2019).

### Conclusion

Creativity theories have long emphasized dual-process models of spontaneous and deliberate thought, but bottom-up data-driven evidence supporting these theories has been largely absent. Using a data-driven eigen-connectivity (EC) analysis with a continuous multitask paradigm (CMP), we extracted latent connectivity patterns shared across multitask FCs - corresponding to deliberate and spontaneous thinking - and we showed that creative cognition may require a balance of these two latent cognitive modes. The EC pattern underlying creative cognition revealed a complex interaction between the two canonical and typically opposite brain networks. We observed creative cognition requires stronger intra-network connectivity in the default mode network (DMN) and fronto-parietal network (FPN), as well as stronger inter-network coupling between the two. We also found higher decoupling in the left-lateralized dorsal attention network (DAN) and visual network, which may facilitate creative thinking by shielding the brain from the external stimuli. In sum, our work provided a bottom-up validation of the latent cognitive modes of creative cognition, offering novel neural evidence for the classic theory of creativity.

## Acknowledgments

This work was supported by a Hasso-Plattner Design Thinking Research Program (HPDTRP) award and an NIH Director’s New Innovator award (MH-119735) to M.S. R.B. was supported by a grant from the National Science Foundation [DRL-1920653].

## Supplemental Materials

**Table S1.** Demographic information.

| Variable | N = 32 |
| --- | --- |
| **Sex** |  |
| Male | 19 |
| Female | 13 |
| **Age** |  |
| Mean ± SD | 30.37 ± 5.41 |
| Range | 19-43 |
| **Handedness** |  |
| Right | 28 |
| Left | 4 |

**Table S2.** Psychometric test results. TTCT: Torrance Test of Creative Thinking; CWIT: Color-Word Interference Test.

| Domain | Psychometric Tests | Mean | Std |
| --- | --- | --- | --- |
| Figural creativity average | TTCT | 119.19 | 22.70 |
| Executive function composite score | CWIT | 115.13 | 14.90 |

**Table S3.** In-scanner behavioral performance.

| Task | Accuracy (mean ± std) |
| --- | --- |
| Theory of Mind | 0.67 ± 0.18 |
| Emotion | 0.96 ± 0.05 |
| VisuoMotor | 0.98 ± 0.04 |
| Working memory | 0.93 ± 0.08 |

**Figure S1.**
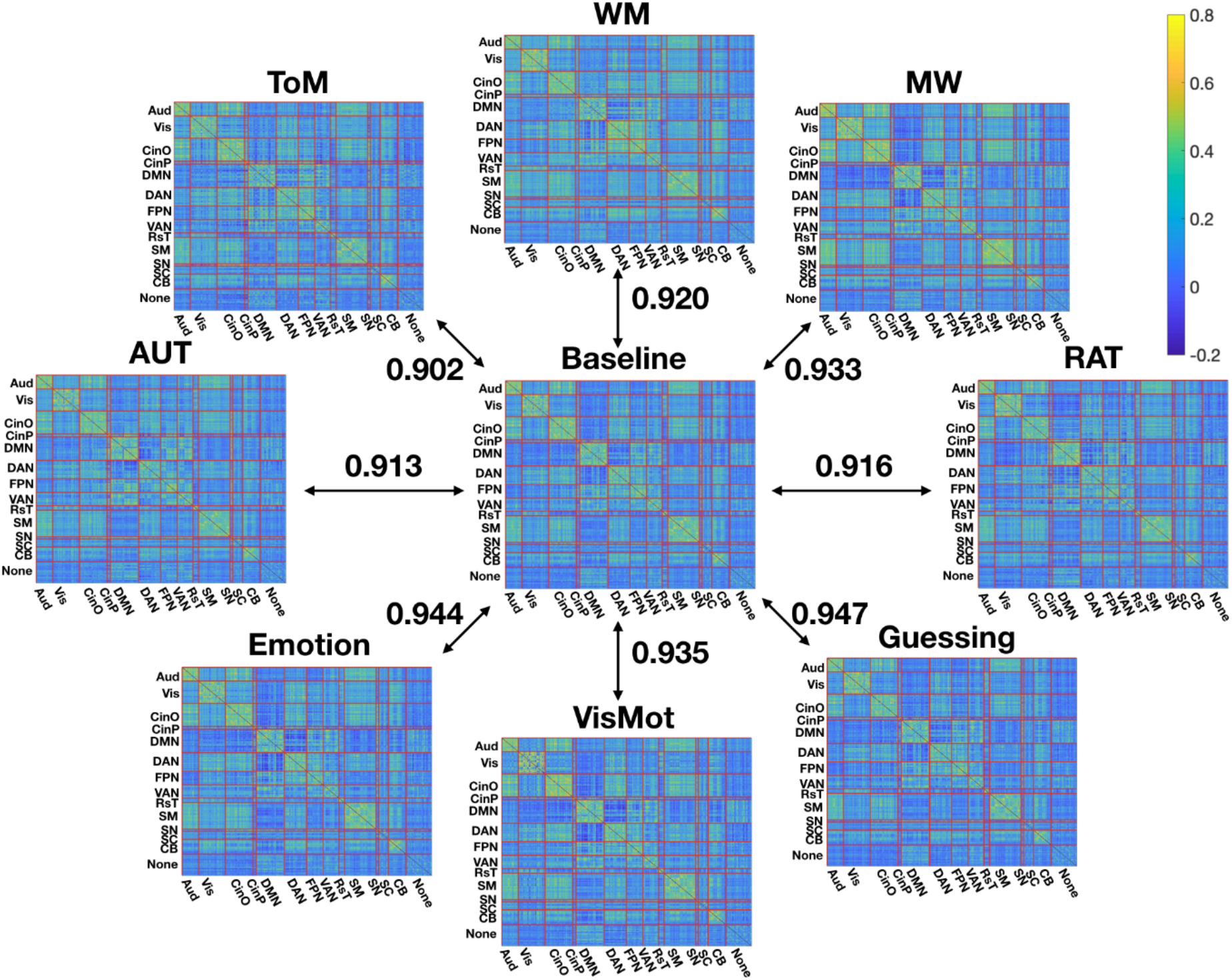
The similarity between group-level task-FCs and baseline-FC was measured by Pearson correlation. Task acronym: WM: working memory; ToM: theory of mind; AUT: alternative uses task; Emotion: emotion task; VisMot: visuomotor task; RAT: remote associates task; MW: mind-wandering. Network acronym: Aud: auditory; Vis: visual; CinO: cingulo-opercular; CinP: cingulo-parietal; DMN: default mode network; DAN: dorsal attention network; FPN: frontal-parietal network; VAN: ventral attention network; RsT: retrosplenial temporal; SM: sensorimotor; SN: salience network; SC: subcortical; CB: cerebellum.

**Figure S2.**
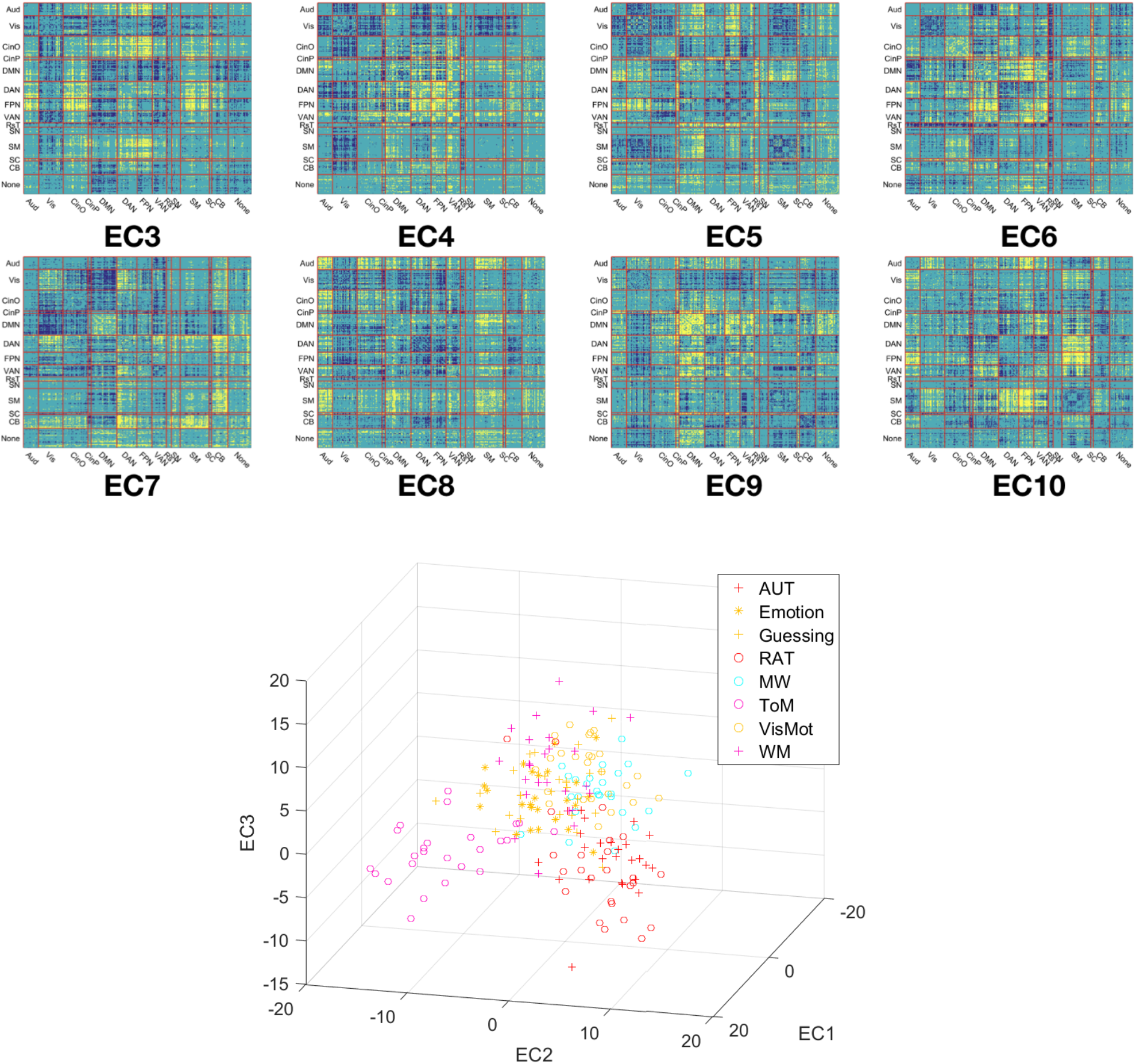
Upper: Thresholded EC3-10 at 10% edge sparsity. Positive edges are shown in yellow and negative edges are shown in dark blue. Lower: Low-dimensional projection using the first three ECs.

**Figure S3.**
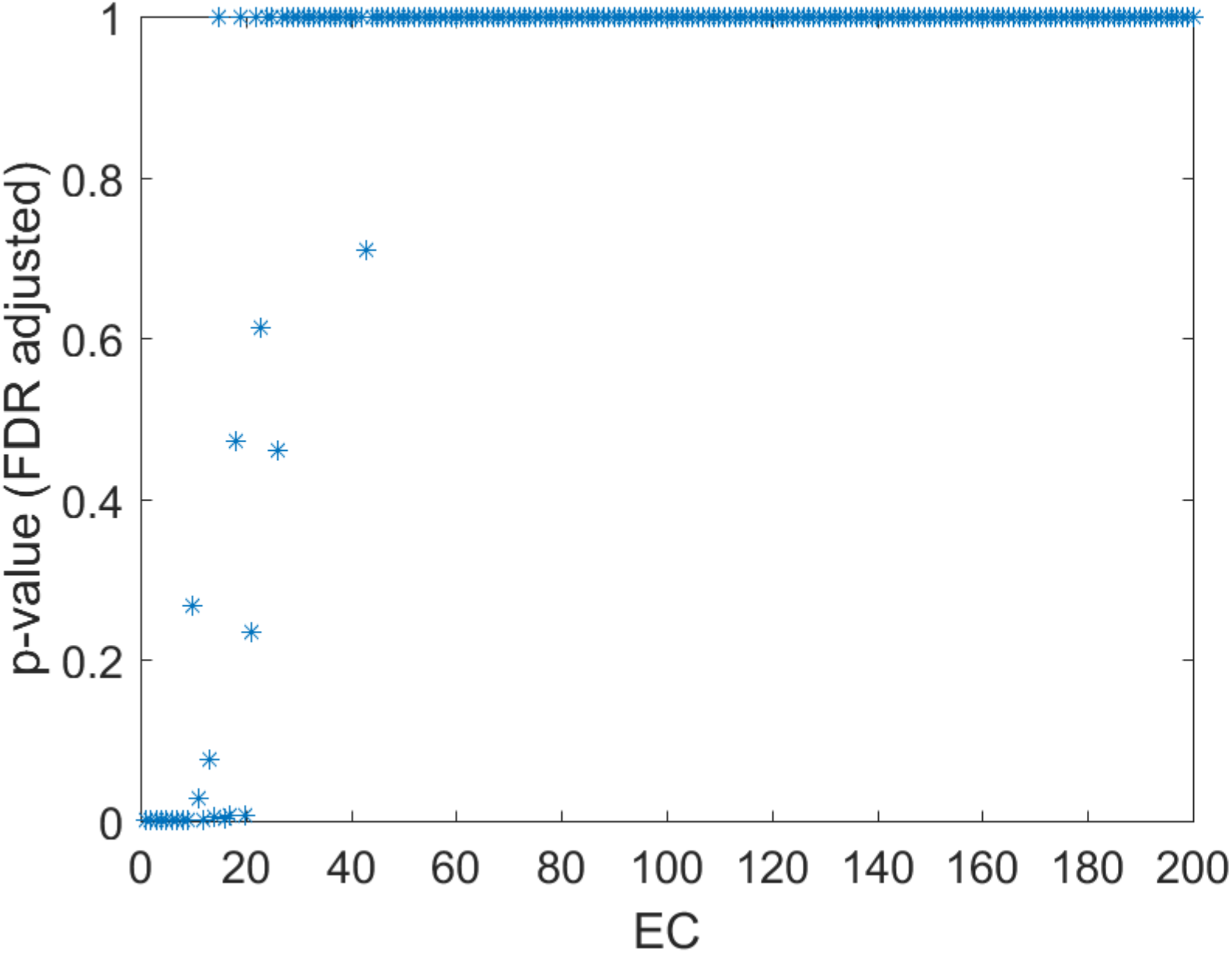
Task separability of EC components. We performed a one-way ANOVA on each EC weight using task labels as grouping variables. Fifteen EC components had significantly different mean across tasks (FDR-corrected *p* < 0.05), suggesting that EC components beyond the first two ECs contained cognitively relevant information.

## Footnotes

1 ToM example: Story: Laura didn’t have time to braid her horse’s mane before going to camp. While she was at camp, William brushed Laura’s horse and braided the horse’s mane for her. Yes/No question: Laura returns assuming that her horse’s hair isn’t braided.

2 RAT example: Cue: dream – break – light. Answer: day.

3 It should be noted that participants spent 40% of the time during the visuomotor task on fixation in between trials, which could have explained VisMot-FCs being projected together with MW.

